# Optimization of 3D bioprinting of human neuroblastoma cells using sodium alginate hydrogel

**DOI:** 10.1101/579268

**Authors:** Jakub Lewicki, Joost Bergman, Caoimhe Kerins, Ola Hermanson

**Affiliations:** Department of Neuroscience, Karolinska Institutet, Biomedicum, SE17177 Stockholm, Sweden

**Keywords:** Bioprinting, Neuroblastoma, Optimization, Alginate, FRESH, POI

## Abstract

There are many parameters in extrusion-based three-dimensional (3D) bioprinting of different materials that require fine-tuning to obtain the optimal print resolution and cell viability. To standardize this process, methods such as parameter optimization index (POI) have been introduced. The POI aims at pinpointing the optimal printing speed and pressure to achieve the highest accuracy keeping theoretical shear stress low. Here we applied the POI to optimize the process of 3D bioprinting human neuroblastoma cell-laden 2% sodium alginate (SA) hydrogel using freeform reversible embedding of suspended hydrogels (FRESH). Our results demonstrate a notable difference between optimal parameters for printing 2% SA with and without cells in the hydrogel. We also detected a significant influence of long-term cell culture on the printed constructs. This observation suggests that the POI has to be evaluated in the perspective of the final application. When taking these conditions into consideration, we could define a set of parameters that resulted in good quality prints maintaining high neuroblastoma cell viability (83% viable cells) during 7 days of cell culture using 2% SA and FRESH bioprinting. These results can be further used to manufacture neuroblastoma *in vitro* 3D culture systems to be used for cancer research.

## 2. Introduction

Stem cell and tumor biology allow for the generation of small organ-like or tumor-like structures to be developed *in vitro*, and this holds great promise for significant improvement of approaches in drug discovery and precision medicine. Bioprinting has emerged as an important tool for improving the conditions and control of such cell culture rationale [1,2]. However, to achieve optimal results, many parameters of the microenvironment must be taken into account. We and others have demonstrated the significance of, e.g., oxygen levels [3,4], substrate stiffness [5,6], substrate roughness [7], and biomaterial properties [8] for progenitor cells to respond properly to external signaling factors such as growth factors, and to execute the appropriate transcriptional programs. Yet, 2-dimensional cell culture is in itself a limiting factor both in stem cell and tumor biology and it has been shown that, for example, certain tumor cells grown in 2D conditions are more sensitive to chemotherapeutic reagents than when grown in three dimensions [9], which may explain some of the lack of progress in cancer research heavily debated during recent years [10,11].

Three-dimensional (3D) bioprinting was first reported to deposit viable cells by Smith et al in 2004 [12]. More than a decade later, 3D bioprinting is constantly being improved in terms of hardware, software, biomaterials, and applications. Standard 3D extrusion-based bioprinters, despite being relatively simple systems, can be challenging to use for cell deposition with high accuracy and viability. Only in recent years, more standardized materials and kits for different applications in bioprinting have become commercially available. Nevertheless, the technology in research settings is still far from a plug-and-play state, mostly due to a high number of variables involved in the 3D bioprinting process.

Some of the key factors are different biomaterials, their concentration and modifications, specific cell types used, cell concentration, deposition process and parameters (for example speed and pressure), crosslinking techniques and parameters, post-processing and cell culture conditions. Before the full potential of the technique can be realized, optimization of these parameters should be performed.

There are several examples of more systematic approaches to the bioprinting optimization process which can serve as a useful entry point for the specific application. However, some of these methods focus mostly or only on printability of the material, not taking the possible biological applications into account [13–15]. Moreover, 3D bioprinting should be viewed through the prism of an additional dimension, namely time. Such constructs may change over time in cell culture conditions due to purely physical interaction and/or dynamics of living cells embedded inside [16,17].

Bioprinting is very often applied for regenerative medicine and tissue engineering research, however, another important filed for this technology is cancer and disease modeling aiming at providing new tools for drug discovery and personalized medicine. Here, we apply a freeform reversible embedding of suspended hydrogels (FRESH) 3D bioprinting method [18] using sodium alginate (SA) for creating constructs populated with human neuroblastoma cells SK-N-BE(2). Neuroblastoma is the most common extracranial childhood tumor that originates from precursor cells in the sympathetic nervous system [19]. There is a number of reports showing that many important physiological aspects of cancer cell culture (including neuroblastoma), such as gene and protein expression, migration and proliferation, are different in 2D cell cultures compared to 3D models [20,21]. Through the creation of more complex and physiologically relevant 3D cancer models, we may gain additional insights into tumorigenesis, progression and treatment.

Given the importance of extracellular matrix (ECM) properties for cell culture and previous observations that stiffer ECM may lead to a reduction of expression of essential transcription factors, such as N-Myc, and also differentiation of neuroblastoma cells [22], we decided to choose SA as a soft hydrogel for cell encapsulation. Moreover, SA has favorable biological and chemical properties, such as low toxicity, nonimmunogenicity, low cost, simple gelation mechanism, and compatibility with 3D bioprinting [18,23,24]. Choosing low concentration SA as a building material presents a challenge for bioprinting process due to its low viscosity. However, a technique such as FRESH could potentially overcome this obstacle. FRESH uses a gelatin slurry for physical support during the printing process and calcium coordination of alginate monomers. To further achieve the highest printing resolution and maximize cell viability, we combined the FRESH approach with an application of the printing optimization index (POI) method [14]. The aim of the POI is to find a set of printing parameters, including a nozzle size, printing speed and pressure, that will result in high accuracy of the printed construct, maintaining low theoretical shear stress (TSS) at the same time. The POI method was originally used with SA and gelatin blends, however, it has previously not been applied to quantitatively assess low concentration SA bioprintability in combination with FRESH technique.

## 3. Materials and methods

### 3.1 Cell culture

SK-N-BE(2) (ATCC, CRL-2271) cells were cultured at 37°C, 5% CO_2_ in DMEM/F12 with GlutaMAX (Life Technologies), supplemented with 10% Fetal Bovine Serum (FBS, Sigma) and 0.1 mg/ml penicillin/streptomycin (Life Technologies). Media was replaced every three to four days. Upon full confluency, cells were passaged 1:5 to uncoated Petri dishes by adding trypLE (Life Technologies) for 5 minutes to dissociate the cells before being resuspended in growth medium and plated.

### 3.2. Sodium alginate preparation

2% SA was prepared by dissolving 20mg/ml SA (Allevi) in SK-N-BE(2) growth media. For the POI assessment 4 mg/ml green fluorescent PLGA microspheres (Sigma) were added to visualize the printed hydrogel during analysis. For printing with cells, SK-N-BE(2) cells were resuspended at 1⋅10^7^/ml of 2% SA.

### 3.3. Control cell encapsulation in 2% SA

SK-N-BE(2) cells were encapsulated in 2% SA at 1⋅10^7^/ml. Encapsulated cells were deposited as 10 μm drops in triplicates using a manual pipette on the bottom of ½ area 96-well optical plate (Corning) and cross-linked using 100 mM CaCl_2_ solution for 15 minutes at 37°C. Subsequently, cross-linker was replaced with fresh SK-N-BE(2) growth medium. After 30 minutes of incubation at 37°C medium was replaced again to reduce the amount of unwashed cross-linker. Then, cells were cultured normally as described above. Live/dead assay (Life Technologies) was performed at 24 and 72h (n=3) and cells were imaged using Operetta CLS high-content screening system (PerkinElmer) using 10x magnification and filters for calcein and EthD detection. Images were then quantified using Harmony 4.5 software (PerkinElmer) using cell detection features.

### 3.4. Gelatin slurry preparation for FRESH 3D bioprinting

Gelatin support gel was prepared using FRESH kit according to the supplier’s manual (Allevi). Briefly, 40 mg/ml of gelatin and 0.16 mg/ml of CaCl_2_ were dissolved in deionized water at 40°C. After overnight incubation at 4°C, glass container with gelatin was filled with cold 0.16 mg/ml of CaCl_2_ and cooled at −20°C until ice crystal formation was apparent. Subsequently, the mixture was blended using the supplied blender (Allevi) in 3 pulses of 30 seconds followed by 30 second breaks between to reduce introduced heat. The resulting blend was centrifuged in 50 ml falcon tubes at 4000 RPM for 2 minutes at 4°C and supernatant was discarded. Gelatin collected at the bottom was resuspended using cold 0.16 mg/ml CaCl_2_ and centrifuged again using the same settings. This step was repeated several times until no white foam was observed on top of the supernatant. Directly before printing, the gelatin slurry was resuspended in cold CaCl_2_ and spun down at 1100 RPM for 5 minutes and the supernatant was discarded. The remaining gelatin was used to fill wells in a 24-well plate. Water-absorbent tissue was laid on top of the wells to draw excess water from the support slurry.

### 3.5. 3D Bioprinting using FRESH method

For each 3D print FRESH method was applied as described previously [18] with some modifications. The Allevi 2 3D bioprinter (Allevi) was used for material deposition in support gelatin slurry using pneumatic extrusion. Each time, a 2.54 cm long 30G blunt needle (Allevi) was used in combination with a 10 ml syringe (BD Biosciences). After printing, 100 mM CaCl_2_ was added to each well containing a scaffold and the plate was placed in the incubator at 37°C and left there for 20 minutes until the gelatin completely dissolved. Afterward, the remaining liquid in each well was replaced with a fresh 100 mM CaCl_2_ pre-warmed to 37°C for further cross-linking at 37°C for 15 minutes. Next, printed constructs were used in different assays described below. When cells were used in 3D bioprints, the cross-linker was replaced with fresh SK-N-BE(2) media. After 30 minutes of incubation at 37°C, medium was replaced again to reduce the amount of unwashed cross-linker.

### 3.6. The POI determination

2% SA mixed with 4mg/ml of green fluorescent PLGA microspheres (Sigma) was printed with or without 1⋅10^7^/ml SK-N-BE(2) cells using the FRESH method described above. One-layer spiral design was used as a blueprint for material deposition. The design was created using Fusion360 software (Autodesk) and processed using Repetier Host (Hot-World GmbH & Co.) to create G-code files. Using Slic3r, the designs were sliced with a line height of 0.2 mm. Printing parameters used for the POI determination are described in the Table 1.

**Table 1.** Printing parameters used for the POI determination.

| Speed<br>(mm/s) | Pressure<br>(psi /<br>kPa) | Number of replicates without cells / **with cells** |  |  |  |
| --- | --- | --- | --- | --- | --- |
|  |  | 5 / **34** | 7.5 / **52** | 10 / **69** | 12.5 / **86** |
| 2 |  | 3 / **3** | 5 / **4** | 5 / **3** | 6 / **3** |
| 4 |  | 4 / **N/A** | 6 / **4** | 6 / **4** | 4 / **4** |
| 6 |  | 4 / **N/A** | 5 / **3** | 6 / **3** | 6 / **4** |
| 8 |  | 3 / **N/A** | 6 / **3** | 6 / **4** | 6 / **4** |

Briefly, the first script changed the analyzed image into binary and then proceeded to the region of interest (ROI) demarcation by a series of dilation and erosion steps resulting in noise reduction. Next, the line was aligned manually, resulting in a solid vertical line. The second script created selection of a one-pixel high box spanning the whole horizontal axis of the image. Within this box, the average width of the line was measured by determining the outmost black pixels and calculating the distance between them. This procedure was run in a loop to analyze the entire length of the imaged line resulting in average line width.

The POI was calculated using equations (1) and (2) as described before [14].

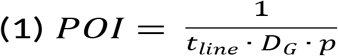

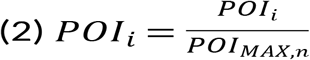

Where D_G_ = needle gauge, p = extrusion pressure, t_line_ = printed line width, POI_MAX_ = the highest POI score found, n = total amount of parameter combinations. POI_i_ score can assume values between 0 (the worst) to 1 (the best).

### 3.7. 3D bioprinting SK-N-BE(2) cells for live/dead assay

For assessing cell viability, SK-N-BE(2) cells were printed at 1⋅10^7^/ml concentration using 7.5 (n=3); 10 (n=3); 12.5 psi (n=4) at 8mm/s. 4-layer lattice G-code file provided by Allevi was used as a blueprint for material extrusion. Scaffolds were cross-linked as described above and cultured for up to 7 days with live/dead assay performed at 24h and 7 days.

Live/dead assay (Life Technologies) was performed according to manufacturer’s instruction using 4 μM EtD and 2 μM calcein. However, DMEM/F12 media was used as washing agent instead of PBS to avoid calcium precipitation and scaffold dissolution. Dead controls were obtained by treating scaffolds with 70% ethanol for 30 minutes at 37°C.

Cells were imaged using LSM 700 confocal microscope (Zeiss) and images were analyzed in 3D using Imaris 9.2 software (Bitplane) using spot detection and surface creation based on the fluorescent signal for later volume calculations.

### 3.8. Statistical analysis

Statistical analysis was performed using Prism 8 (GraphPad). For line width measurements, two-way ANOVA and Tukey’s multiple comparison test were used. For SK-N-BE(2) cells viability after printing, two-way ANOVA with Sidak’s multiple test were performed. Object volumes were compared using one-way ANOVA with Tukey’s multiple comparison test. Results were considered significant at p ≤ 0.05. For correlation between pressure and viability of cells at 24h, Pearson correlation was calculated.

## 4. Results

### 4.1. SK-N-BE(2) cells encapsulation in 2% SA

There have been several reports of using alginates for neuroblastoma cell growth, however, either focusing on mouse cell lines and/or peptide-modified alginates [26–28]. Therefore before proceeding with bioprinting, simple SK-N-BE(2) cell encapsulation in 2% SA gel followed by viability assay was performed to asses biocompatibility with the human neuroblastoma cell line. After 24 hours post-encapsulation SK-N-BE(2) cells displayed 56% of viability and which remained stable at the later time point at 72 hours (Fig. 1). Cells appeared round in morphology, homogenously filling the entire volume of casted SA gel in a well plate. After 72 hours in culture, cells covered more volume of the gel and started to form larger colonies (Fig. 1a). This positive initial result, proved SA to support human neuroblastoma cell viability and growth upon encapsulation, making it a possible candidate for bioprinting application.

**Figure 1.**
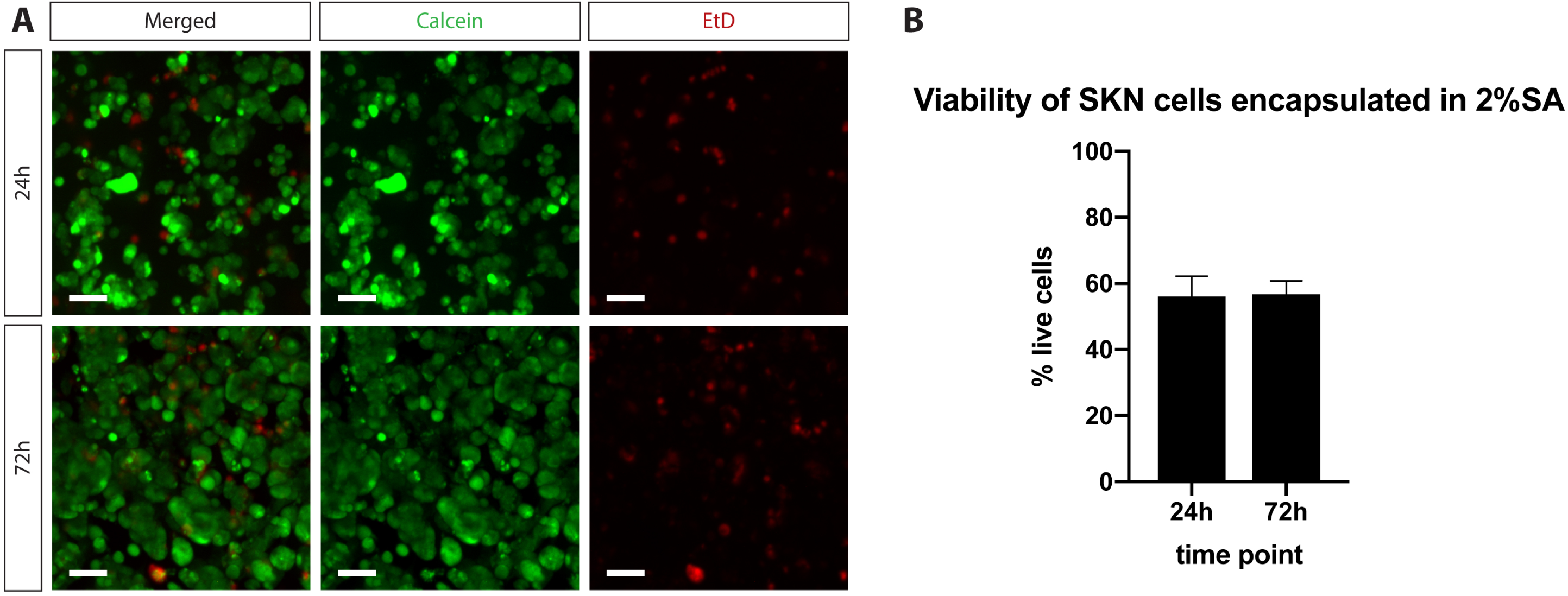
The viability of SK-N-BE(2) cells upon encapsulation in 2% SA. (A) Fluorescent microscopy images of live/dead assay performed on encapsulated SK-N-BE(2) cells at 24 and 72h. Green (calcein) indicates live cells, red (EthD) indicates dead cells. Scale bars represent 50 μm. (B) Quantification of live/dead assay. Bars show mean % of cell viability + SD.

### 4.2. Parameter optimization index

First, 2% SA alone was used for the POI determination (POI_2% SA_). Simple single-layer spiral design was used as a template for material extrusion. Four different speeds and four different extrusion pressures were used to deposit SA in a support gelatin bath (Table 1.).

Accurate width measurement of a printed strand is central for the POI determination, thus we developed an ImageJ macro to evaluate this parameter in semi-automated and non-biased approach. This script allowed us to measure extrusion width along the entire line, pixel-by-pixel, resulting in an accurate average strand dimension that was later used for POI.

As shown in Figure 2, the image of a printed SA line after crosslinking acquired with a fluorescent microscope is later converted to a binary image which is used to create a region of interest (ROI). Proper alignment of an automatically generated ROI with both brightfield and fluorescent microscopic image (Fig. 2e) proved this method to be a fast and efficient way to analyze SA prints.

**Figure 2.**
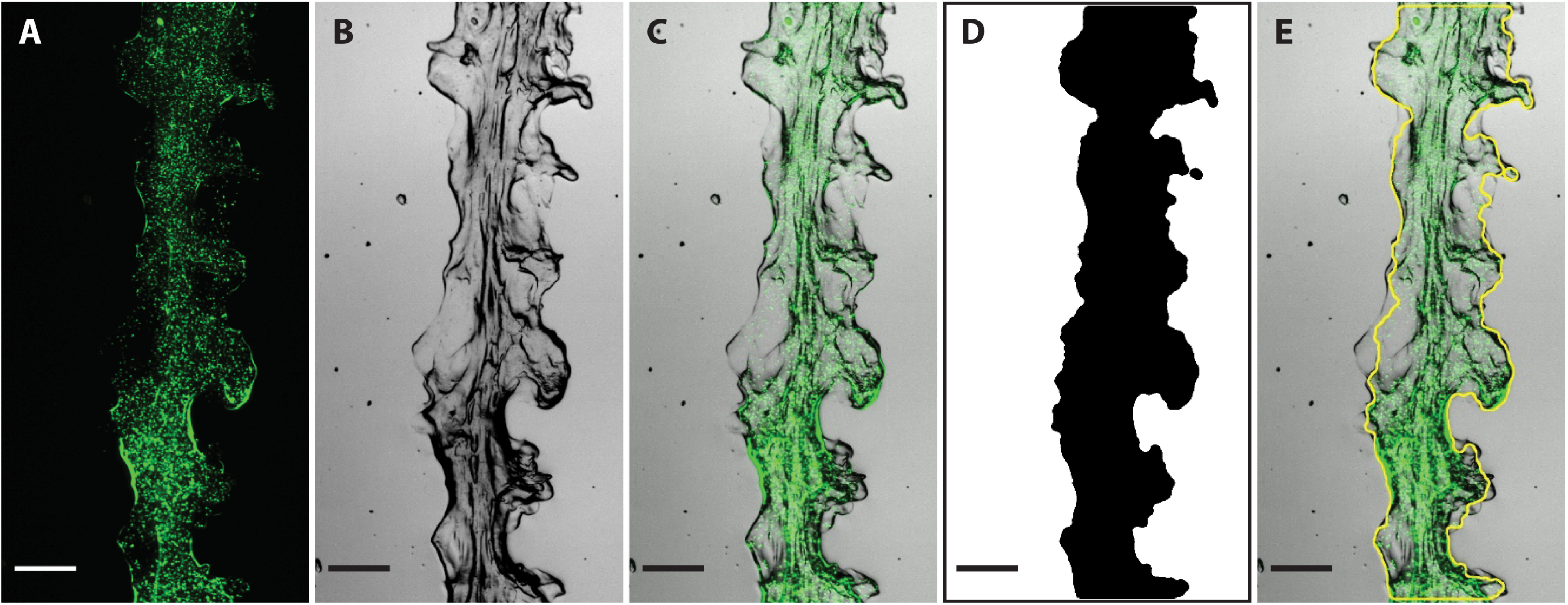
Printed line width analysis. (A) Fluorescent microscopy image of an example line printed using 2% SA and the FRESH method. Green signal comes from green fluorescent PLGA microspheres for visualization. (B) The same line imaged using phase-contrast microscopy. (C) Fluorescent and phase-contrast images merged together. (D) A binary representation of the line as a result of image processing using custom ImageJ script. (E) Merged image from (C) with yellow line overlaid on the top showing region of interest (ROI) based on binary line representation. ROI is finally used for line width measurement. Scale bars indicate 200 μm.

Analysis of printed lines (Fig. 3a) using only 2% SA showed that pressure is the most important factor influencing strand width. 72.8% of variability between different groups could be accounted to pressure, whereas speed was responsible for only 2.3% of the total variation between groups. The thinnest lines were achieved while printing with 5 psi (291 to 302 μm of average width depending on speed), whereas the thickest were a result of using the highest tested pressure: 12.5 psi (557-584 μm of average width depending on speed).

**Figure 3.**
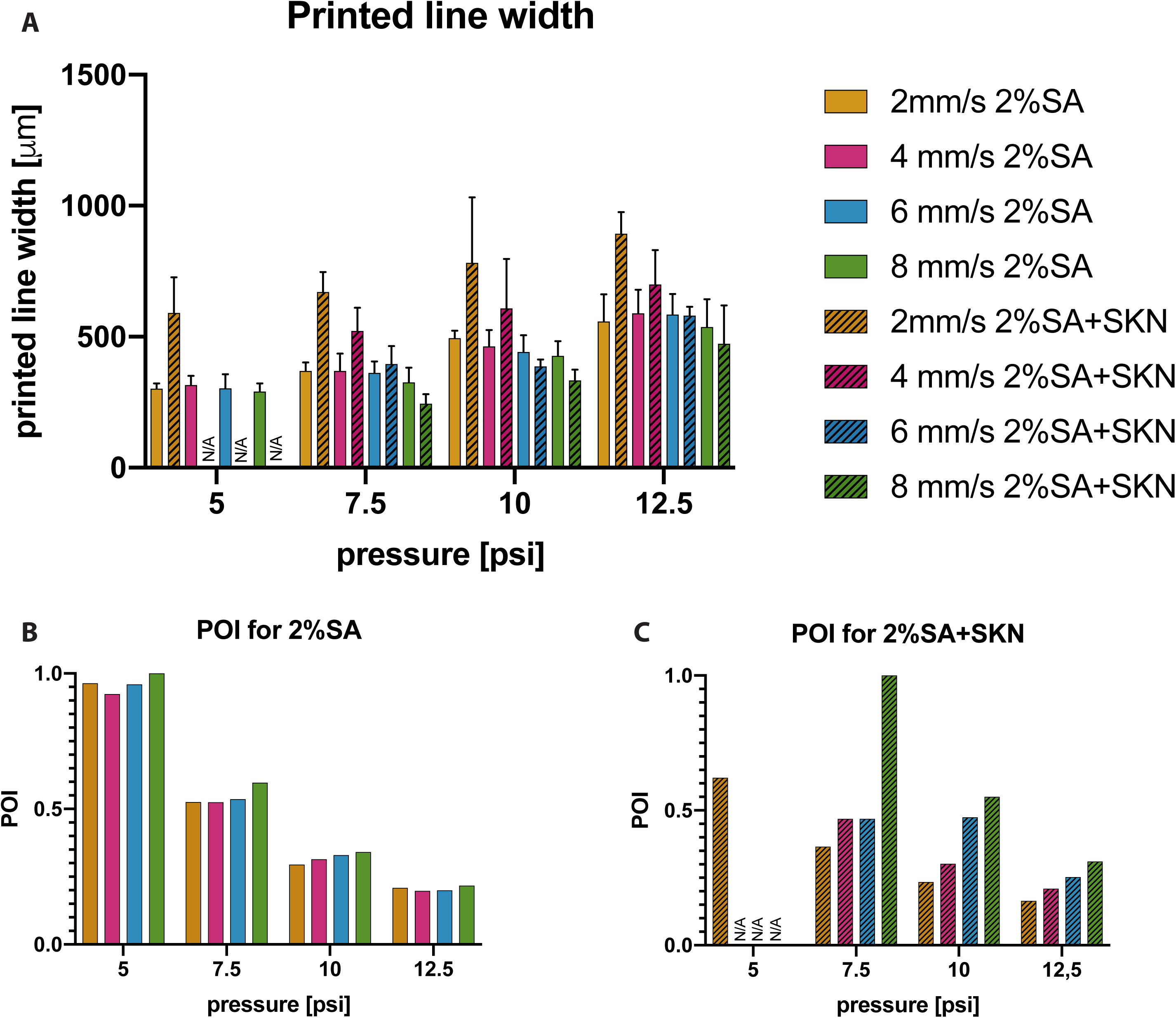
Parameter optimization index for 2% SA with and without cells. (A) Printed line width quantification. 2% SA with SK-N-BE(2) cells and without them was printed using a set of different pressures and speeds. Bars represent mean line width quantified with custom ImageJ script + SD. Results of Tukey’s multiple comparison test are presented in Supplementary Figure 1. (B) the POI for 2% SA without cells. (C) the POI for 2% SA with 1⋅10^7^/ml SK-N-BE(2) cells. Values were calculated based on equation (1) and bars represent normalized POI values from equation (2).

Subsequently, these values were used to calculate POI_2%SA_. Maximum normalized POI_2%SA_ value of 1 was a result of using combination of the lowest pressure (5 psi) and highest speed (8 mm/s) providing parameters with the best print accuracy and the lowest TSS for printing cells (Fig 3b). However, when we used this set of parameters to print 2% SA mixed together with SK-N-BE(2) cells, we observed very poor mechanical properties of the final constructs, resulting in a quick structural disintegration during handling. Thus, we decided to repeat printed line width measurements, this time using 2% SA and SK-N-BE(2) cells combined together.

Indeed, during our analysis we were unable to obtain enough intact samples for strand dimensions analysis while using 5 psi at 4, 6 and 8 mm/s printing speed. Additionally, the width of the printed lines using 2% SA with SK-N-BE(2) cells was significantly different from lines printed with 2% SA alone when using the lowest speed at every tested pressure (Supplementary Figure 1.). Within constructs populated with cells, not only pressure was a significant factor accounting for 18.0% of total variance, but also speed (50.3% of total variance).

A new POI was calculated using measurements from printing 2% SA with SK-N-BE(2) cells (POI_2%SA+SKN_). Maximum POI_2%SA+SKN_ was obtained for 7.5 psi and 8mm/s printing speed (Fig 3c). Altogether, these results revealed the impact of the presence of cells in the tested material and its influence on the POI determination. Hence the parameters calculated for 3D printing of pure biomaterial samples might not be suitable for the same biomaterial when mixed with cells.

### 4.3. FRESH bioprinting of SK-N-BE(2) cells

Next, we used 2% SA mixed with SK-N-BE(2) cells to FRESH bioprint a four-layer lattice (Fig. 4). 7.5, 10, and 12.5 psi pressures were used at 8 mm/s. 5 psi pressure was not used due to poor mechanical properties of the prints. The highest speed from the previous tests was applied, as this resulted in the highest POI scores in each pressure condition (Fig. 3c). Such constructs were cultured for up to one week and viability assay was performed at 24h and 7 days.

**Figure 4.**
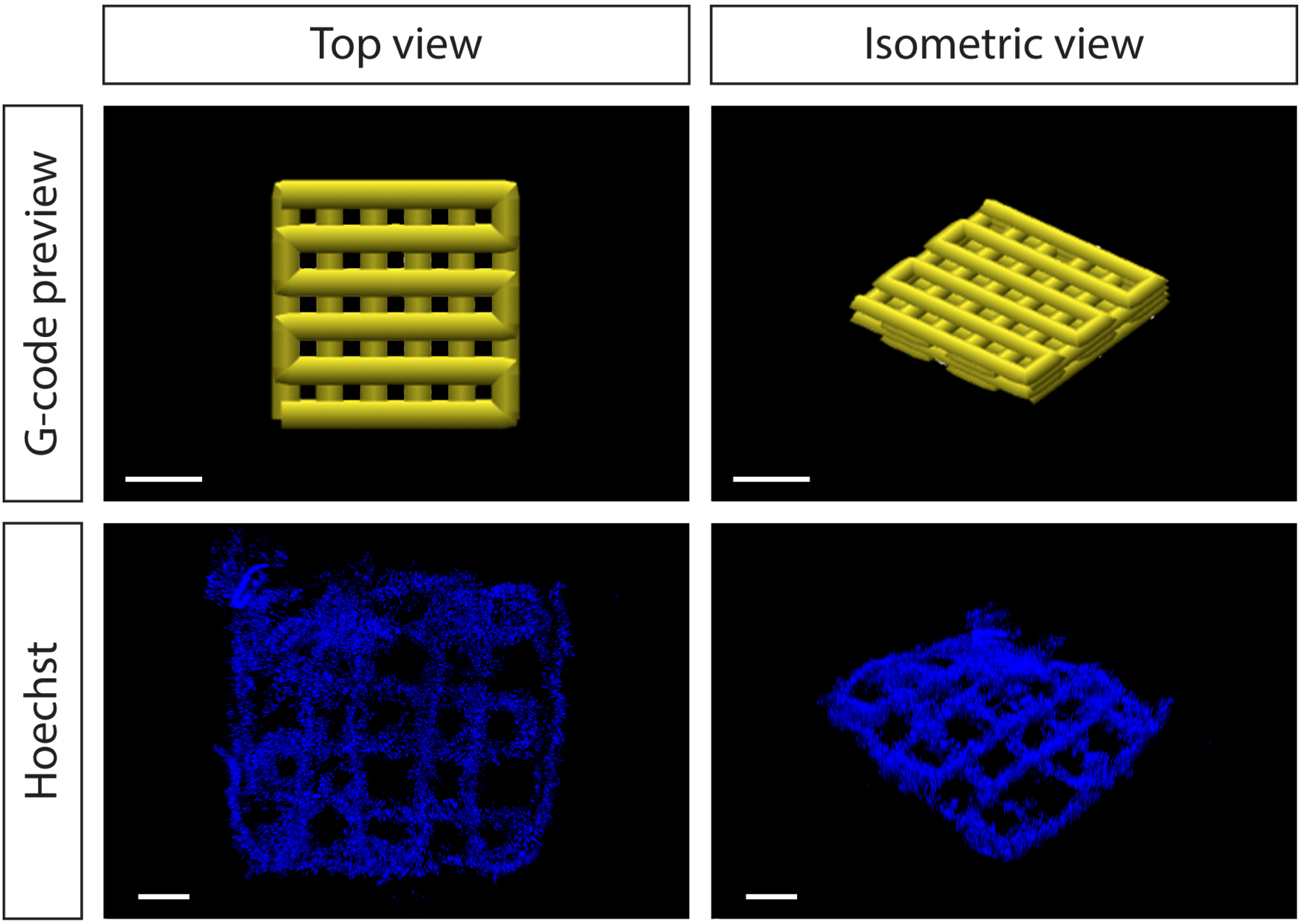
Geometry used for FRESH bioprinting of 2% SA with SK-N-BE(2) cells. 4-layer lattice is represented as a G-code visualization on the top panel. The bottom panel shows confocal microscopy images of 2% SA with 1⋅10^7^/ml SK-N-BE(2) cells extruded at 12.5 psi and 8 mm/s 24h after printing. Blue signal represents Hoechst nuclei staining of the cells. Scale bars represent 1 mm.

24 hours post-printing cells were homogenously distributed in the entire print volume. At this time point cells displayed relatively low viability. There was a significant difference in cell survival while using different pressures with a strong positive correlation between the pressure applied and the percentage of live cells (R^2^ = 0.99). At 24h post-printing only 19% of cells were viable at 7.5 psi, however, it was more than doubled at 10 psi (40% of live cells) and 52.5% viability for the highest pressure (Fig. 5a, b).

**Figure 5.**
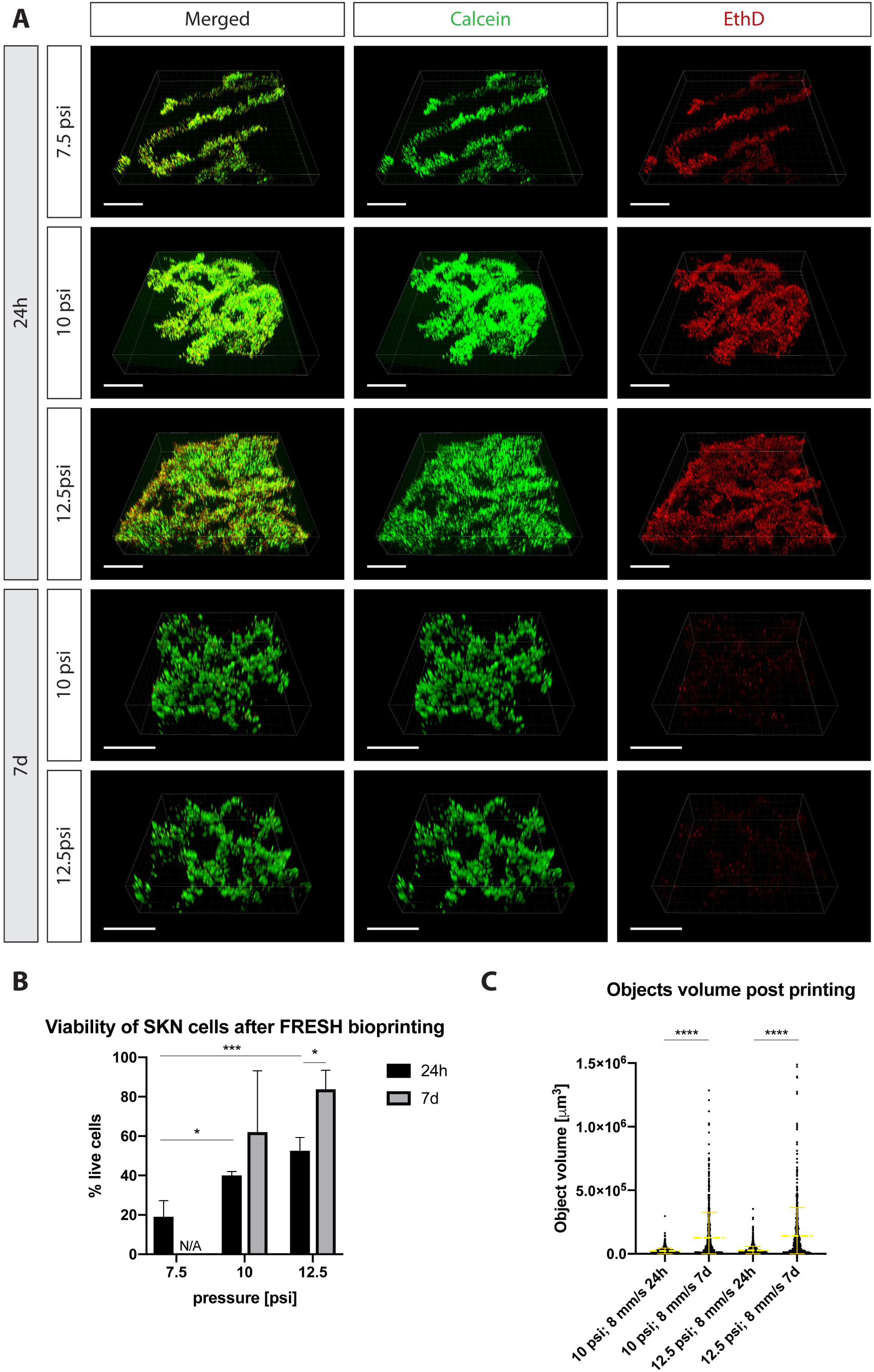
SK-N-BE(2) cells viability after printing using 2% SA and FRESH. (A) Confocal microscopy images of the live/dead assay performed at 24 and 7d after printing with a different set of extrusion parameters. Green (calcein) indicates live cells, red (EthD) indicates dead cells. Scale bar represents 1 mm. (B) Quantification of live/dead assay. Bars represent mean % of cell viability + SD. (C) Quantification of object volumes from live/dead assay on SK-N-BE(2) cells after printing. Scatter plot shows separate data points representing single volumes of objects detected based on calcein cytoplasmic staining and image segmentation using Imaris software. Asterisks over plots refer to statistical significance in multiple comparison test. * P≤ 0.05; *** P ≤ 0.001; **** P ≤ 0.0001.

The post-printing cell viability after 7 days was notably higher with 62% at 10 psi and reaching significant difference at 83.7% viability at 12.5 psi (p = 0.03). Constructs printed with 7.5 psi were more fragile than others, and did not withstand the staining process for viability assays (data not shown). Cells over time started to form more compact colonies and clustered together. Volume analysis of such clusters, showed significantly higher mean volume of clusters at 7 days compared to 24h: 1.3⋅10^5^ μm^3^ for 10 psi and 1.4⋅10^5^ μm^3^ for 12.5 psi at 7d; 2.3⋅10^4^ μm^3^ and 2.8⋅10^4^ μm^3^ for 10 and 12.5 psi at 24h respectively.

Together, our results show that despite low initial viability, SK-N-BE(2) cells are able to recover and display significantly higher viability at later time points. We further suggest using a POI value corrected for long time cell culture effect on the construct. Maximum POI_2%SA+SKN_ did not result in the robust constructs after culturing for 7 days. However, printing with parameters for the next highest POI_2%SA+SKN_ values (10 and 12.5 psi at 8mm/s) gave rise to scaffolds that survived culturing and post-processing, maintaining high cell viability.

## 5. Discussion

The material chosen here, sodium alginate, is widely used for cells encapsulation both *in vitro* [29–31] and *in vivo* [32,33]. Even though unmodified alginate-based hydrogel does not support interaction with cells directly [34] it is possible to use them as a scaffold for cell immobilization that can lead to aggregation [35]. However, if needed, alginate can be modified with specific cell attachment proteins or blended with different biomaterials such as silk fibroin to improve cell adhesion [36,37]. Lack of cell adhesion sites in polysaccharide chains of sodium alginate may explain to some extent initial lower viability of SKN cells encapsulated in it [38]. Further, reduction of cell viability upon bioprinting can be a combination of shear stress on extruded cells and cross-linking conditions that are slightly different in the FRESH printed samples comparing to simply casted hydrogel [18,39]. During the FRESH bioprinting, cells need to pass through a long and thin canal of a needle (2.54 cm long and 0.159 mm inner diameter), whereas in our encapsulation control, cells mixed with 2% SA were dispensed using standard pipette tip with an inner nozzle diameter of around 1.5 mm. This radically different geometry will result in higher shear stress upon bioprinting. Additionally, exposure to CaCl_2_ as a cross-linker can reduce cell viability by generating osmotic stress and/or apoptosis induction through calcium signaling [40,41]. During FRESH bioprinting, cells are exposed to calcium ions for a longer time then in encapsulation due to time need for printing and gelatin support dissolving, which can further explain differences in cell viability between these two conditions.

Another aspect is a difference in cell survival dependent on the pressure applied for extrusion. There are studies on this relationship showing that increased pressure results in decreased cell viability due to mechanical stress introduced [39,42]. However, here we show a reverse relationship. Cells displayed the highest viability in the highest pressure applied (12.5 psi) at 24h and 7 days post-printing. This contradictive results could be accounted for the fact that pressures that we studied were generally on the lower end of the scale (from 5 to 12.5 psi), whereas other reports mentioned above describe differences in cell viability only for bigger changes in pressure (i.e. 5 vs 20 psi). Significant differences in cell survival presented here are probably result of a different mechanism that comes into play and is stronger than mechanical stress due to cell extrusion. This, for example, could be an effect of geometry (thinner lines extruded with lower pressure) or result of different overall cell number deposited at a single construct, but more studies would be required to explain the exact mechanism. Nevertheless, 7 days after bioprinting, SK-N-BE(2) cells displayed much higher viability. This could be due to washing away dead cells from the construct, cell recovery from mechanical damage, and/or increased proliferation. Large cell clusters observed at this time point could support the cell proliferation effect also observed for MC3T3-E1 cells in alginate [43].

The success of particular bioprinting application relays on the optimization of all the components in the given system. There are reports on optimization of specific aspects of bioprinting, from biocompatibility of materials used [44], to extrusion process [13], however combination of different aspects or long term effects are sometimes overlooked. The POI may serve as a valuable tool for maximizing print resolution with control of TSS, but it is important to note, that these calculations should be made not on the biomaterial alone, but in combination with target cell type at the desired concentration. The presence of cells can alter the rheological properties of the hydrogel changing parameters required for proper extrusion [45]. Furthermore, it can also have a long-term effect on the structure itself. Proliferation, migration, and scaffold remodeling can affect mechanical properties, shape, and integrity of the construct [17]. Also, the cell culture conditions, for example, the presence of ions such as Na^+^ or Mg^2+^ may lead to calcium release from the alginate gel and its eventual dissolution [46]. Thus, in summary, to create reproducible and useful bioprinted *in vitro* models, it is important to take all these factors into account.

## 6. Conclusion

We applied the POI method to find the best settings for printing SK-N-BE(2) cells embedded in a 2% SA hydrogel. We showed the importance of using the POI analysis on the final composition of the biomaterial printed, including the right concentration of cells, as it significantly affected the outcome. Printing neuroblastoma cells with parameters for the highest POI_2%SA+SKN_ (7.5 psi and 8 mm/s) despite being a reflection of the best printing accuracy and the lowest TSS, resulted in fragile constructs that did not stand staining process. However, applying speed and pressure from the next highest POI_2%SA+SKN_ values (10 and 12.5 psi at 8mm/s) resulted in constructs with high cell viability after 7 days in cell culture. Therefore, we suggest using the POI as a tool for finding optimal parameters in the context of the final application. If the bioprinted construct is intended to be used at later time points, the POI should be viewed in the perspective of long-term cell culture and its effects on cell viability and scaffold integrity.

## Acknowledgements

We thank members of the Hermanson lab for valuable input on the manuscript. This study was supported by project grants from the Swedish Research Council (VR-MH), the Swedish Cancer Society (Cancerfonden), and the Swedish Childhood Cancer Foundation (Barncancerfonden) to O.H.

## Abbreviations

FRESH: freeform reversible embedding of suspended hydrogels
POI: parameter optimization index
SA: sodium alginate

## SUPPLEMENTARY FILES

**Supplementary Figure 1.**
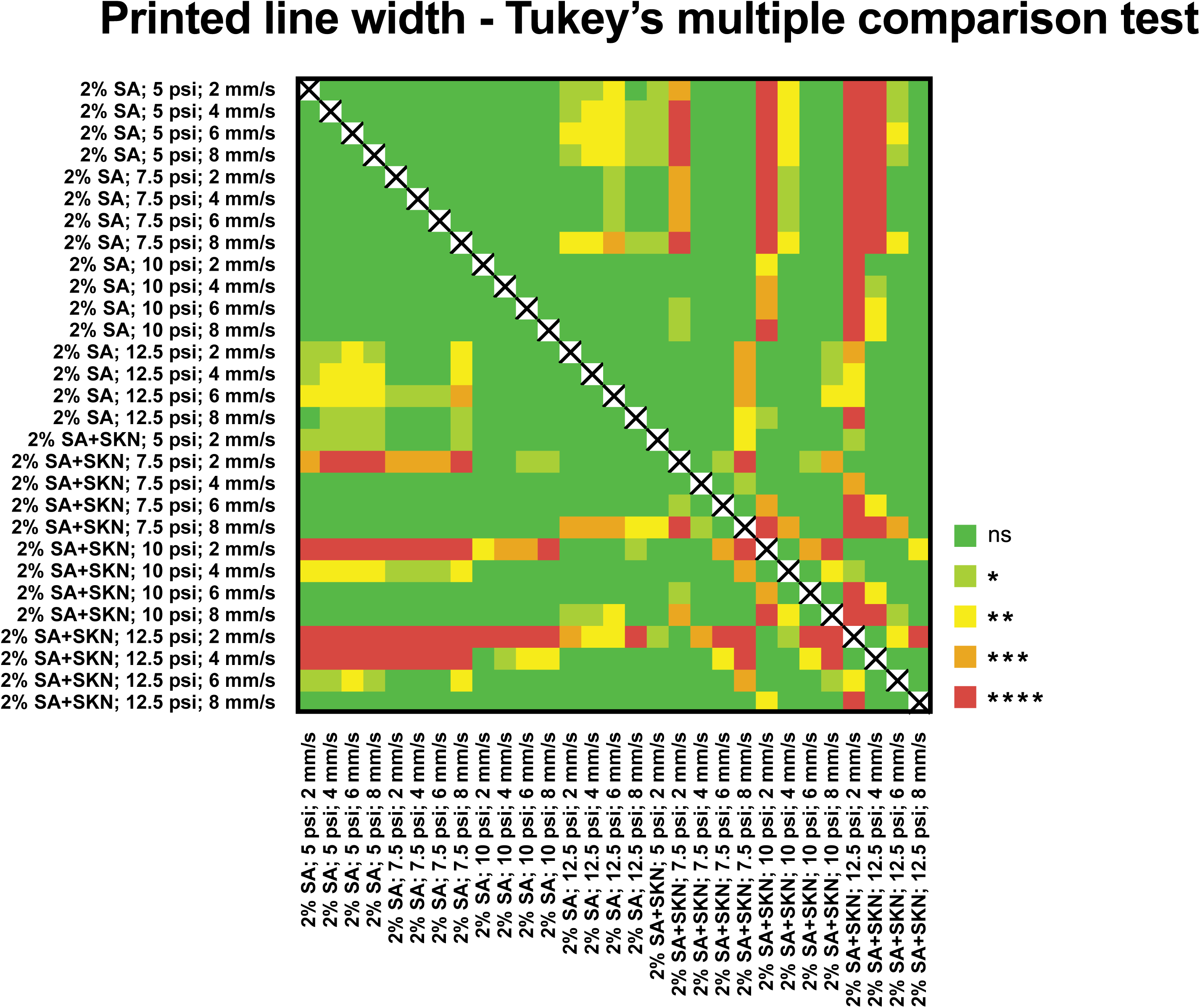
Results of Tukey’s multiple comparison test for average width of lines printed with 2% SA with or without SK-N-BE(2) cells. Ns P > 0.05; * P ≤ 0.05; ** P ≤ 0.01; *** P ≤ 0.001; **** P ≤ 0.0001.

